# Peripheral modulation of Pumilio in intestinal stem cells and the *corpus allatum* affects sleep latency in *Drosophila*

**DOI:** 10.1101/2025.11.09.687465

**Authors:** Josué A. Rodríguez-Cordero, Marialena Dorta-Avilés, Imilce A. Rodriguez-Fernandez, Alfredo Ghezzi, José E. Lizardi-Ortiz, José L. Agosto-Rivera

## Abstract

While central circuits governing sleep are well-studied, the contribution of signaling from peripheral tissues remains a critical yet less understood aspect of sleep regulation. The highly conserved RNA-binding protein Pumilio (Pum) is a post-transcriptional regulator expressed in multiple tissues that influence systemic physiology, but its role in modulating basal sleep has not been established. Although Pumilio’s function in central neurons has been linked to sleep homeostasis following deprivation, whether it regulates sleep through peripheral mechanisms remains unknown. Here, we use conditional genetic tools in the fruit fly *Drosophila melanogaster* to demonstrate that Pumilio acts in the intestinal stem cells (ISCs) and the endocrine *corpus allatum* (CA) to specifically regulate the transition to sleep. Reducing Pumilio function in either the ISCs or the CA independently and significantly accelerates nighttime sleep onset, while overexpression produces the opposite effect. This behavioral change is accompanied by widespread transcriptional alterations in the brain, characterized by a robust upregulation of genes involved in cellular stress responses, including Heat shock protein 83 (*Hsp83*). Our findings reveal a previously unrecognized gut-endocrine-brain signaling axis and identify peripheral post-transcriptional regulation as a key input to the central control of sleep behavior.

## Introduction

Sleep is a fundamental and highly conserved biological process essential for physiological homeostasis and optimal cognitive function. Difficulties in initiating sleep, a condition known as sleep-onset insomnia, are a primary symptom of insomnia disorders. These conditions represent a significant clinical challenge, affecting approximately 10% of the adult population worldwide and underscoring the importance of understanding their distinct regulatory mechanisms [1–4]. Current pharmacological interventions for insomnia, which include GABA receptor and melatonin receptor agonists, mainly target the central nervous system [5]. While sleep is primarily governed by central brain circuits, contributions from peripheral tissues are increasingly recognized as critical determinants of behavioral state [6–9].

Systemic signals originating from organs, such as the gut, communicate the body’s metabolic, microbial, and immunological status to the brain through hormones and metabolites, linking internal physiology to complex behaviors, including appetite, eating behavior, and sleep [10–12]. Understanding this inter-organ communication may therefore be key to dissecting how distinct aspects of sleep, such as its initiation and maintenance, are differentially regulated. The fruit fly, *Drosophila melanogaster*, with its robust genetic toolkit and conserved sleep-like state, serves as a genetically amenable model for dissecting how specific peripheral signaling pathways impinge upon central circuits to modulate these facets of sleep [13–15].

Pumilio (Pum) is a highly conserved RNA-binding protein that mainly functions as a post-transcriptional repressor. First identified for its critical role in establishing abdominal patterning in the *Drosophila* embryo, Pumilio achieves its function by binding to specific sequences, most commonly in the 3^*′*^ untranslated regions of target mRNAs [16]. Often in complex with cofactors such as Nanos and Brat, Pumilio orchestrates the translational repression or stabilization of transcripts, thereby fine-tuning protein expression [17–19].

In recent years, the gut–brain axis has emerged as an important bidirectional communication network linking intestinal physiology with neural activity and behavior. Growing evidence indicates that gut-derived signals, shaped by epithelial renewal, microbial composition, and immune status, influence brain function and sleep–wake cycles through circulating metabolites, hormones, and inflammatory mediators [20, 21]. Within this context, intestinal stem cells (ISCs) play a pivotal role in maintaining epithelial integrity and responding to physiological stress [22]. Moreover, studies in both flies and mice have shown that sleep loss disrupts gut homeostasis and ISC function [23, 24], positioning ISCs as potential modulators of systemic signals that impact brain function and behavioral states such as sleep initiation.

In this study we performed conditional, tissue-specific genetic manipulations to uncover a novel role for Pumilio in regulating sleep latency. Our results demonstrate that reducing Pumilio function in two distinct peripheral sites, intestinal stem cells and the endocrine *corpus allatum* (CA), significantly and independently accelerates the onset of nighttime sleep. This discovery suggests a coordinated or parallel control system originating from metabolically and hormonally active tissues. To gain molecular insights into the brain’s response to these peripheral signals, we performed transcriptomic analysis on heads of flies in which *pumilio* was knocked down in the ISCs and CA. This analysis revealed that peripheral *pumilio* manipulation induces widespread gene expression changes in the brain, most notably a robust upregulation of genes involved in cellular stress responses. Together, these results define a novel gut- and endocrine gland-to-brain signaling network that specifically regulates the transition to sleep, highlighting post-transcriptional regulation in the periphery as a critical input for central behavioral control.

## Materials and methods

### snRNAseq data from Fly Cell Atlas

The t-SNE plots for *pumilio* single-nucleus expression in adult intestinal tissue were obtained from ASAP (https://flycellatlas.org/asap), using publicly available snRNA-seq data from the Fly Cell Atlas [25] (GEO accession GSE120537). To visualize normalized expression by cell annotation, the corresponding loom file was downloaded from ASAP, accessed using loompy [26] (v3.0.8), and plotted with seaborn [27] (v0.13.2) using Python (v3.12).

### Fly stocks

*Drosophila melanogaster* stocks were reared on Nutri-Fly Bloomington Formulation media (Genesee Scientific, San Diego CA) under a 12:12 hr light/dark cycle. Flies for TARGET system [28] experiments were raised at 18°C; all other stocks were raised at 25°C.

The following stocks were obtained from the Bloomington Drosophila Stock Center (BDSC) unless otherwise noted: UAS-pum^RNAi^ (VALIUM10, RRID: BDSC 26725), UAS-pum^RNAi^ (VALIUM20, RRID: BDSC 36676), UAS-GFP (VALIUM10, RRID: BDSC 35786), UAS-mCherry^RNAi^ (VALIUM20, RRID: BDSC 35785), UAS-EGFP^RNAi^ (VALIUM22, RRID: BDSC 41558). These RNAi lines were generated by the TRiP project [29]. We also used UAS-Pum (provided by Michael Stern (Rice University) and described by Schweers et al. [30])), UAS-tubGal80^ts^ (RRID: BDSC 7017). UAS-tdTomato (RRID: BDSC 36328), UAS-RpL3-FLAG (RRID: BDSC 77132), Aug21-Gal4 (RRID: BDSC 30137), ISC-KCKT^ts^-GAL4 (RRID: BDSC 91411), esg-Gal4, UAS-2xYFP; Su(H)GBE-Gal80, tub-Gal80^ts^ (prepared by S. Hou (NIH, USA) provided by Heinrich Jasper lab (Buck Institute for Research on Aging, now at Genentech Inc.)). Within the article “ts” is used as shorthand for tubGal80^ts^.

See Table A in S1 File for genotypes used in each experiment.

### Sleep assays

Flies were allowed to mate for two days post-eclosion and were transferred to fresh food every 2-3 days until they reached the experimental age of 6-8 days. For all experiments, the age difference between individuals was no more than two days.

For sleep assays, individual flies were briefly anesthetized with CO_2_ and placed in 5 mm polycarbonate tubes containing either standard 5% sucrose in 2% agar or Nutri-Fly media as specified. The tubes were placed in *Drosophila* Activity Monitors (DAMs; TriKinetics, Waltham, MA) inside an environmentally controlled incubator set to 80% humidity and a 12:12 hr light/dark cycle. Data were acquired using the DAMSystem3 software (TriKinetics). DAM data was analyzed using MATLAB.

Following an initial acclimation period, baseline activity was recorded for several days. For experiments utilizing the TARGET system, transgene expression was controlled by shifting the temperature [28]. The TARGET system allows for temporal and spatial control of gene expression by combining the *Gal4*-UAS binary system [31] with a ubiquitously expressed, temperature-sensitive version of the *Gal4* repressor, *Gal80* (*tubGal80*^*ts*^). At the permissive temperature (18°C), *Gal80*^*ts*^ is active and inhibits *Gal4*, preventing it from driving expression of the UAS-linked transgene. When flies are shifted to the restrictive temperature (29°C), *Gal80*^*ts*^ is inactivated, allowing *Gal4* to function and induce transgene expression. Sleep was defined as any period of five or more consecutive minutes of inactivity [13, 32]. Nighttime sleep latency was calculated as the time from lights-off to the first bout of sleep [33].

Sleep assay statistical analyses were performed using GraphPad Prism (Version 10.5.0, GraphPad Software, La Jolla, CA). For comparisons between two experimental groups on a given night, differences were assessed using the non-parametric Mann-Whitney test, unless otherwise stated. Data from experiments with multiple groups across several days, such as the RNAi control and dietary challenge experiments, were analyzed using a mixed-effects model with a Geisser-Greenhouse correction, followed by Tukey’s post-hoc test for multiple comparisons. All data are presented as mean *±* s.e.m.

### RNAseq

Following the sleep assay, flies of each genotype were pooled and allowed to recover for 30 minutes. Flies were then flash-frozen in liquid nitrogen. Heads were separated from bodies by repeated vortexing and freezing, followed by separation through a sieve. For each biological replicate, 10 heads were pooled in 1 mL of TRI Reagent (Sigma-Aldrich) and homogenized with RNAse-free plastic pestles.

Total RNA extraction and sequencing was outsourced to Novogene (Sacramento, CA, USA) using the company’s standard procedures. Stranded mRNA sequencing libraries were prepared from purified poly-A-containing mRNA and sequenced on an Illumina NovaSeq platform to generate approximately 12 Gb of 150 bp paired-end reads per sample.

For bioinformatic analysis, raw reads were first assessed for quality using FastQC v0.12.1 [34] and trimmed for adapters and low-quality sequences using Trimmomatic v0.39 [35]. Trimmed reads were then aligned to the *Drosophila melanogaster* reference genome (dm6) using STAR 2.7.11b [36]. Gene expression was quantified, and differential expression analysis was performed using DESeq2 [37]. Gene Ontology (GO) term and pathway enrichment analysis were conducted using GeneCodis4 [38].

All RNA-seq data generated in this study will be made publicly available through the NCBI Gene Expression Omnibus (GEO) under accession number [to be provided upon acceptance].

### Imaging

Widefield fluorescence microscopy images were acquired on a Nikon Eclipse microscope equipped with a Nikon DS-Qi2 camera and controlled by NIS-Elements BR 4.30.01 software. A 10x objective was used for image capture, utilizing a standard red fluorescence filter. Images were recorded as 1608 x 1608 pixel frames with 1×1 binning, an exposure time of 1 second, and a gain of 1.0x. The resulting image calibration was 0.73 µm/pixel. Images were cropped to focus on the head and neck of the files.

## Results

### Intestinal expression of *pumilio*

To characterize the expression pattern of *pumilio* (*pum*) within the adult *Drosophila* intestine, we analyzed publicly available single-nucleus RNA sequencing (snRNA-seq) data from the Fly Cell Atlas [25] (GEO accession GSE120537) (Fig 1a). The analysis revealed that *pum* is broadly expressed across diverse intestinal cell types (Fig 1a and b).

**Fig 1.**
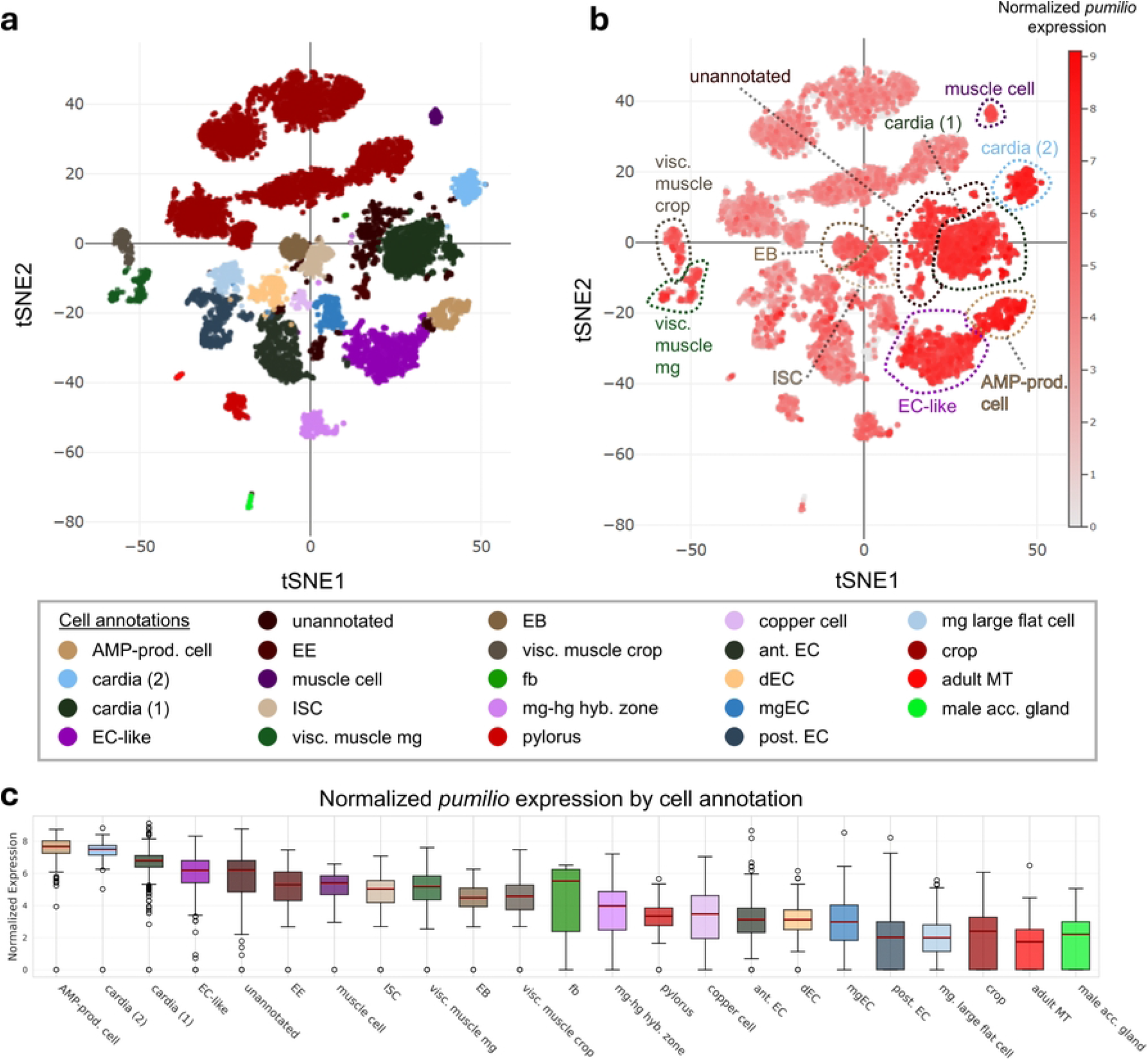
Expression of *pumilio* across adult *Drosophila* intestinal and associated cell types. (a) tSNE plot of single-nucleus RNA-seq data from the adult *Drosophila* gut showing annotated cell clusters from the Fly Cell Atlas [25] (GEO accession GSE120537). Legend for cell annotation using different colors per cluster. (b) Normalized *pumilio* expression projected onto the same t-SNE embedding, visualized using ASAP (https://flycellatlas.org/asap). Red intensity indicates log-normalized expression values (Seurat). Dashed outlines highlight clusters with high *pumilio* expression. (c) Boxplots showing normalized *pumilio* expression across annotated cell categories. Solid red line marks the median. *Abbreviations*: adult MT, adult Malpighian tubule; AMP-prod. cell, antimicrobial peptide-producing cell; ant. EC, anterior enterocyte; cardia (1) and cardia (2), distinct cardia cell clusters; dEC, differentiating enterocyte; EB, enteroblast; EC-like, enterocyte-like cell; EE, enteroendocrine cell; fb, fat body; ISCs, intestinal stem cells; mgEC, midgut enterocyte; mg-hg hyb. zone, midgut–hindgut hybrid zone; mg large flat cell, midgut large flat cell; muscle mg, midgut visceral muscle; visc. muscle crop, visceral muscle of the crop; post. EC, posterior enterocyte; pylorus, pyloric region; copper cell, copper cell region; crop, crop epithelium; male acc. gland, male accessory gland.

Notably, *pum* showed robust expression within the key progenitor and signaling lineages of the gut, including intestinal stem cells (ISCs), the enteroblasts (EBs), and the hormone-secreting enteroendocrine (EE) cells (Fig 1c). While the highest expression levels were detected in specialized populations such as antimicrobial peptide-producing (ECs) and cardia cells (forming the proventriculus), the significant expression in the ISC lineage suggested a potential role for *pum* in regulating intestinal renewal and inter-organ communication. This observation motivated subsequent experiments aimed at determining the functional consequences of manipulating Pumilio levels within ISCs.

### Conditional knockdown of *pumilio* in *esg+ Su(H)GBE-Gal80* –restricted cells (ISCs and CA) accelerates sleep onset in male flies

Based on the expression of *pumilio* in ISCs, we next sought to determine its functional role in these cells using conditional RNA interference (RNAi) in *Drosophila melanogaster* via the TARGET system (*Gal4/Gal80*^*ts*^) [28, 31]. We used the *esg-Gal4* driver, which is widely used to target ISCs and EBs since ISCs were first characterized in 2006 [39, 40]. However, it has recently been reported that this driver also directs expression in the endocrine *corpus allatum* (CA) [41]. We therefore assessed the effects of manipulating *pum* expression in both ISCs and the CA.

To restrict expression to ISCs while excluding EBs, we used a fly line carrying *esg-Gal4* combined with *Su(H)GBE-Gal80* transgenes. In addition, the line carries a transgene encoding a temperature-sensitive *Gal80* repressor (*tub-Gal80*^*ts*^) driven by the ubiquitous tubulin promoter, which allows temporal control of *Gal4* activity. This TARGET line was crossed to different UAS transgenes to modulate *pumilio* expression under temperature control. Experimental flies (*esgGal4*^*ts*^ *>UAS-pum*^*RNAi*^) and controls (*esgGal4*^*ts*^ *>UAS-GFP*) were placed in *Drosophila* Activity Monitors and maintained at 18°C, a temperature at which transgene expression does not occur, for a 60-hour baseline period, and then shifted to 29°C to induce *pum*^*RNAi*^ or GFP expression in ISCs and the CA. This combination restricts *esg-Gal4* expression to intestinal stem cells while excluding enteroblasts, and also drives expression in the *corpus allatum*.

Sleep profiles (Fig 2a) and representative raster plots (Fig 2b) were generated for each genotype under the different temperature regimes, and distinct features of sleep architecture were analyzed (Fig 2c-h). Induction of *pum*^*RNAi*^ at 29°C significantly reduced nighttime sleep latency in male flies relative to controls (Fig 2d). Although total nighttime sleep was also significantly altered (Fig 2c), the reduction in sleep latency was the most consistent and pronounced effect of *pum* knockdown. Other sleep architecture parameters, such as episode number and duration, showed no overall differences (Fig 2e-h).

**Fig 2.**
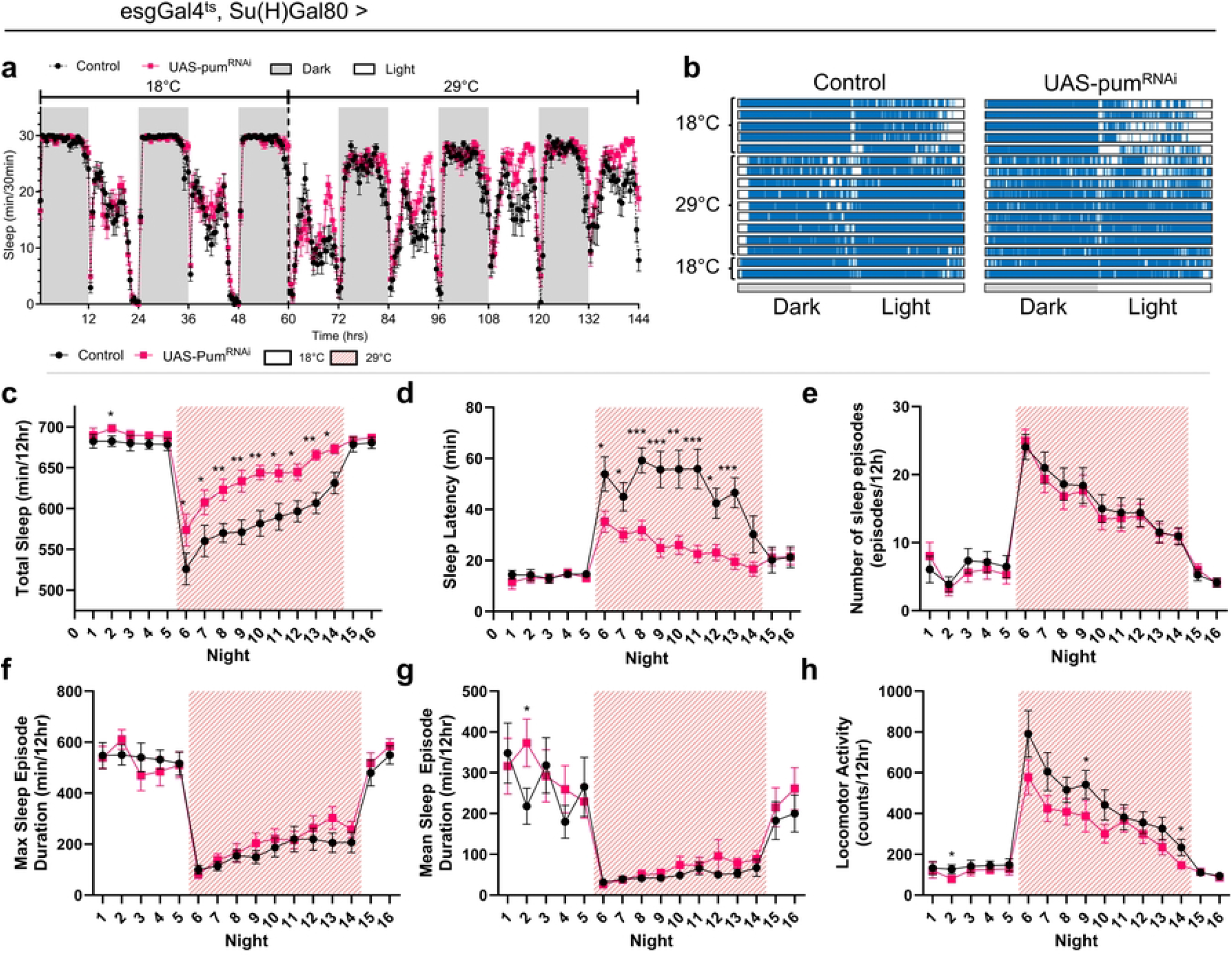
Conditional knockdown of *pumilio* in ISCs and CA reduces nighttime sleep latency in adult males. (a) Sleep profile comparing *pum*^*RNAi*^ male flies (*esgGal4*^*ts*^, *Su(H)Gal80 >UAS-pum*^*RNAi*^ (VALIUM10); pink line) and control flies (*esgGal4*^*ts*^, *Su(H)Gal80 >UAS-GFP*; black line), showing average sleep (min/30 min). Data represents 60 hours of baseline period at 18°C, followed by 84 hours with transgene expression at 29°C. Light and dark cycles are indicated by white and grey bars, respectively. (b) Raster plots of representative individual control and *pum*^*RNAi*^ male flies showing their 24-hour sleep patterns during baseline (18°C) and knockdown expression (29°C) periods. Each row represents one day; blue indicates sleep (≥ 5 min inactivity). (c-h) Quantification of nighttime sleep parameters over 16 consecutive nights. Flies were maintained at 18°C for 5 nights (baseline), shifted to 29°C for 9 nights (induced knockdown, indicated by shaded background), and returned to 18°C for 2 nights (reversal). For (a) and (c-h), values are mean ± s.e.m (* *P <*0.05, ** *P <*0.01, *** *P <*0.001, Mann-Whitney test). Genotypes: Control (*esgGal4*^*ts*^, *Su(H)Gal80 >UAS-GFP*, n=15), *pum*^*RNAi*^ (*esgGal4*^*ts*^, *Su(H)Gal80 >UAS-pum*^*RNAi*^ (VALIUM10), n=17).

Similar trends in sleep latency reduction upon *pumilio* knockdown were observed in female flies (Figure A in S1 File). The core phenotype of reduced sleep latency in males was reproducible across biological replicates (Figure B in S1 File). While both sexes exhibited changes in sleep latency, the effect was consistently stronger and more reproducible in males. Therefore, subsequent analyses focused on male flies to facilitate robust comparisons and minimize variability across experiments.

Importantly, this phenotype was specific to *pumilio* inactivation, as demonstrated by comparisons with control RNAi constructs (Figure C and Table B in S1 File), and was dependent on *Gal4*-mediated expression, with no effects observed from leaky transgene expression in the absence of the driver (Figure D in S1 File). Furthermore, the sleep latency phenotype persisted regardless of dietary conditions, as both standard Nutrifly food (Bloomington recipe) and 5% sucrose alone yielded the same *pum*^*RNAi*^ sleep phenotypes (Figure E and Table C in S1 File).

### Conditional overexpression of *pumilio* in *esg+ Su(H)GBE-Gal80* –restricted cells (ISCs and CA) tends to delay sleep onset in male flies

To determine whether overexpressing *pumilio* levels had an opposing effect, we performed conditional overexpression of *pumilio* using the same temperature-sensitive TARGET system (*esgGal4*^*ts*^, *Su(H)Gal80 >UAS-pum*). Male flies overexpressing *pumilio* were compared to genetic controls lacking either the UAS or *Gal4* transgene. Upon temperature shift to 29°C, *pumilio* overexpression led to an increase in nighttime sleep latency compared to control flies carrying the overexpression construct without the *Gal4* driver or a UAS-flag-tagged control construct (Fig 3). This phenotype, opposite to that observed upon *pum* knockdown, further supports a role for Pumilio in modulating the timing of sleep initiation.

**Fig 3.**
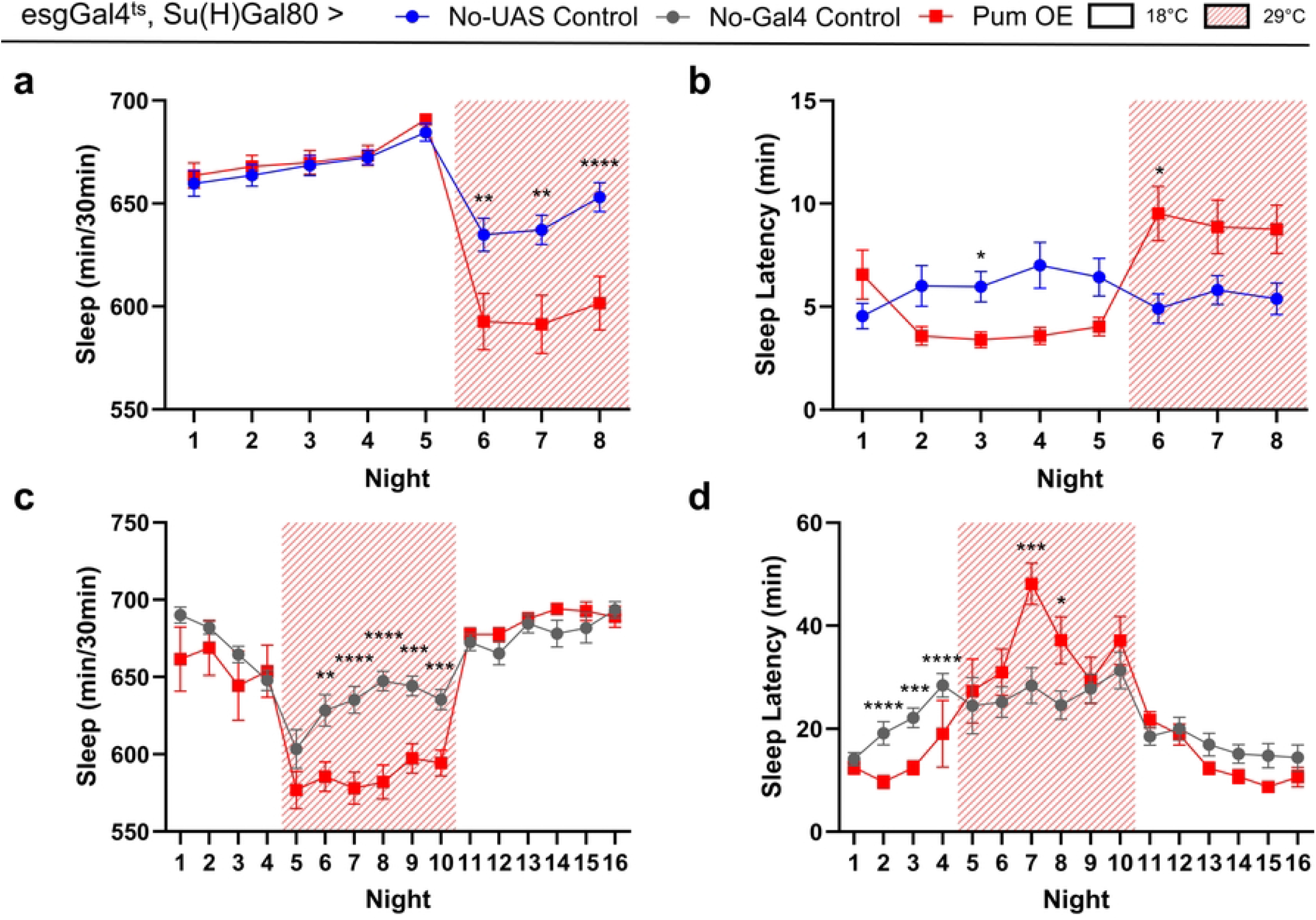
Conditional overexpression of *pumilio* in ISCs and CA tends to increase nighttime sleep latency in male flies. (a,b) Nighttime total sleep (a) and sleep latency (b) over an 8-night period. Experimental flies (Pum OE; *esgGal4*^*ts*^ */ +*; *Su(H)Gal80 / UAS-pum, UAS-RpL3-Flag*, n = 64) were compared to control flies lacking the UAS-*pum* transgene (No-UAS Control; *esgGal4*^*ts*^ */ +*; *Su(H)Gal80 / UAS-RpL3-Flag*, n = 63). (c,d) Nighttime total sleep (c) and sleep latency (d) over a 16-night period. Experimental flies (Pum OE; *esgGal4*^*ts*^, *Su(H)Gal80 >UAS-pum*, n = were compared to a driverless UAS control (No-Gal4 Control; *CyO / +; UAS-pum / +*, n = 28). For all panels, temperature was shifted from 18°C to 29°C (indicated by shaded regions) at the beginning of night 6 to induce overexpression. Values and error bars are mean *±* s.e.m. (* *P <*0.05, ** *P <*0.01, *** *P <*0.001 for comparisons by Mann-Whitney test).

### Head transcriptome analysis reveals systemic transcriptional responses to peripheral *pumilio* knockdown

To investigate the molecular changes in the brain that might underlie the observed sleep phenotype following *pum* knockdown in *esg+ Su(H)GBE-Gal80* –restricted cells, we performed RNA sequencing (RNAseq) on heads from *pum* kd (*esgGal4*^*ts*^; *Su(H)Gal80 >UAS-pum*^*RNAi*^) and control (*esgGal4*^*ts*^; *Su(H)Gal80 >mCherry*^*RNAi*^) male flies. This approach enabled the identification of transcriptional responses in the central nervous system, potentially driven by systemic signals from the gut or CA. Behavioral validation confirmed reduced sleep latency in the cohort used for RNAseq (Fig 4a).

**Fig 4.**
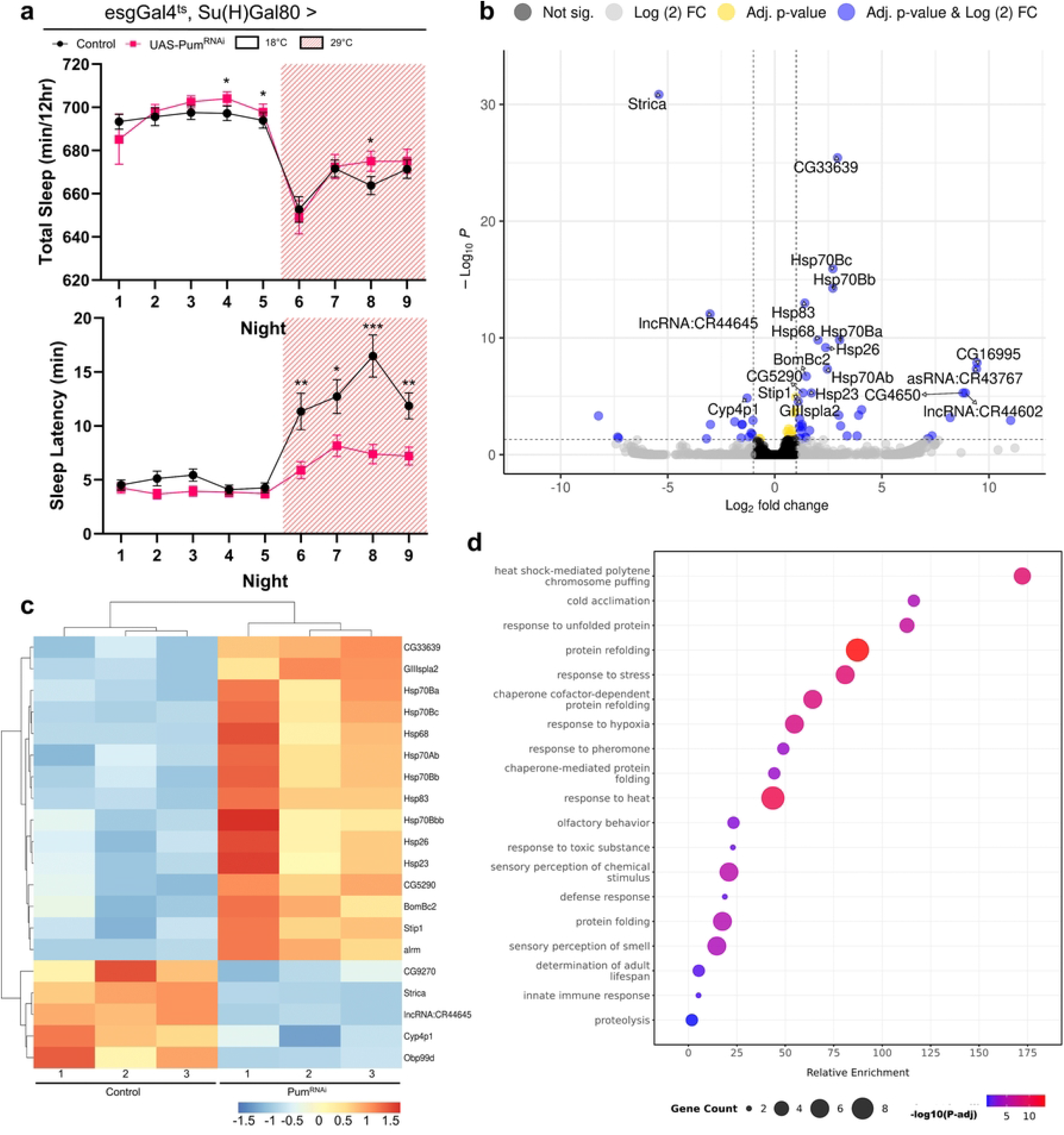
RNAseq of *Drosophila* heads reveals transcriptome-level changes related to altered sleep latency after *pumilio* knockdown in ISCs and CA. (a) Nighttime total sleep (left) and sleep latency (right) for male flies used in RNAseq experiments, comparing control (*esgGal4*^*ts*^, *Su(H)Gal80 >UAS-mCherry*^*RNAi*^, n = 63) with *pum* knockdown flies (*esgGal4*^*ts*^, *Su(H)Gal80 >UAS-pum*^*RNAi*^ (VALIUM20), n = 62). *Pumilio* knockdown flies show significantly lower sleep latency, while no overall difference in total sleep was observed. Values are mean ± s.e.m. Asterisks indicate statistically significant differences (* *P <*0.05, ** *P <*0.01, *** *P <*0.001, Mann-Whitney test). (b) Volcano plot illustrating differentially expressed genes (DEGs) from the RNAseq analysis. Significantly up- and down-regulated genes are highlighted in blue (*P*-adj *<*0.05, |log_2_FoldChange| ≥ 1), with key genes labeled. (c) Heatmap displaying hierarchical clustering of the top 20 (lowest *P*-adj DEGs). Cutoffs for differentially expressed genes were *P*-adj *<*0.05 and |log_2_FoldChange|*≥* 1. (d) Gene Ontology (GO) enrichment analysis of the 55 DEGs. −log_10_(*P*-adj) shown as a gradient of blue to red. Multiple enriched clusters correspond to temperature and stress responses.

Differential expression analysis identified 55 genes significantly altered upon *pum* knockdown (*P*-adj *<*0.05, |log_2_FoldChange|≥ 1), as visualized in the volcano plot (Fig 4b; full list of differentially expressed genes Table D in S1 File; full DESeq2 results in S2 File). A heatmap of the top 20 differentially expressed genes (DEGs) illustrates distinct transcriptomic profiles between *UAS-pum*^*RNAi*^ and control flies, demonstrating clustering of samples by genotype (Fig 4c).

Gene Ontology (GO) analysis of the 55 DEGs revealed significant enrichment for several functional categories (Fig 4d). Notably, terms related to “response to heat”, “response to unfolded protein”, and “protein folding” were prominent, consistent with the strong upregulation of multiple heat shock protein genes. Among these, *Hsp83* was one of the most significantly upregulated DEGs (log_2_FoldChange = 1.40, *P*-adj = 1.04E-13) and appears in the top 20 DEGs shown in the heatmap. Other enriched GO categories included “defense response”, “innate immune response”, “response to hypoxia”, and “olfactory behavior”. The DEGs contributing to these categories included several odorant-binding proteins (e.g., *Obp19a, Obp28a, Obp56a, Obp56h, Obp99d*), genes involved in lipid metabolism (e.g., *Lsd-1, Acsx2*), cytochrome P450s (e.g., *Cyp4p1, Cyp4e3*), *NT5E-2* (encoding an ecto-5^*′*^-nucleotidase), and various non-coding RNAs (e.g., *lncRNA:CR44645, asRNA:CR43767*), indicating broad physiological impacts of peripheral *pumilio* depletion on the head transcriptome.

### Pumilio function in both ISCs and the CA contributes to sleep latency control

Given that the *esgGal4; Su(H)Gal80* transgenes can drive *Gal4* in both ISCs and the CA [41], we next aimed to determine the relative contributions of these cells and tissues to the sleep latency phenotype. We performed targeted *pum* knockdown using more specific drivers: *ISC-KCKT*^*ts*^*-GAL4* for ISCs [42] and *Aug21-Gal4*^*ts*^ for the CA [43, 44].

Targeted *pum* knockdown in ISCs significantly reduced nighttime sleep latency compared to controls (Fig 5a), with no concurrent change in total sleep duration. Likewise, CA-specific *pum* knockdown also led to a significant decrease in nighttime sleep latency (Fig 5b), without affecting total sleep. These results indicate that pumilio activity in both ISCs and the CA independently contributes to the regulation of nighttime sleep onset.

**Fig 5.**
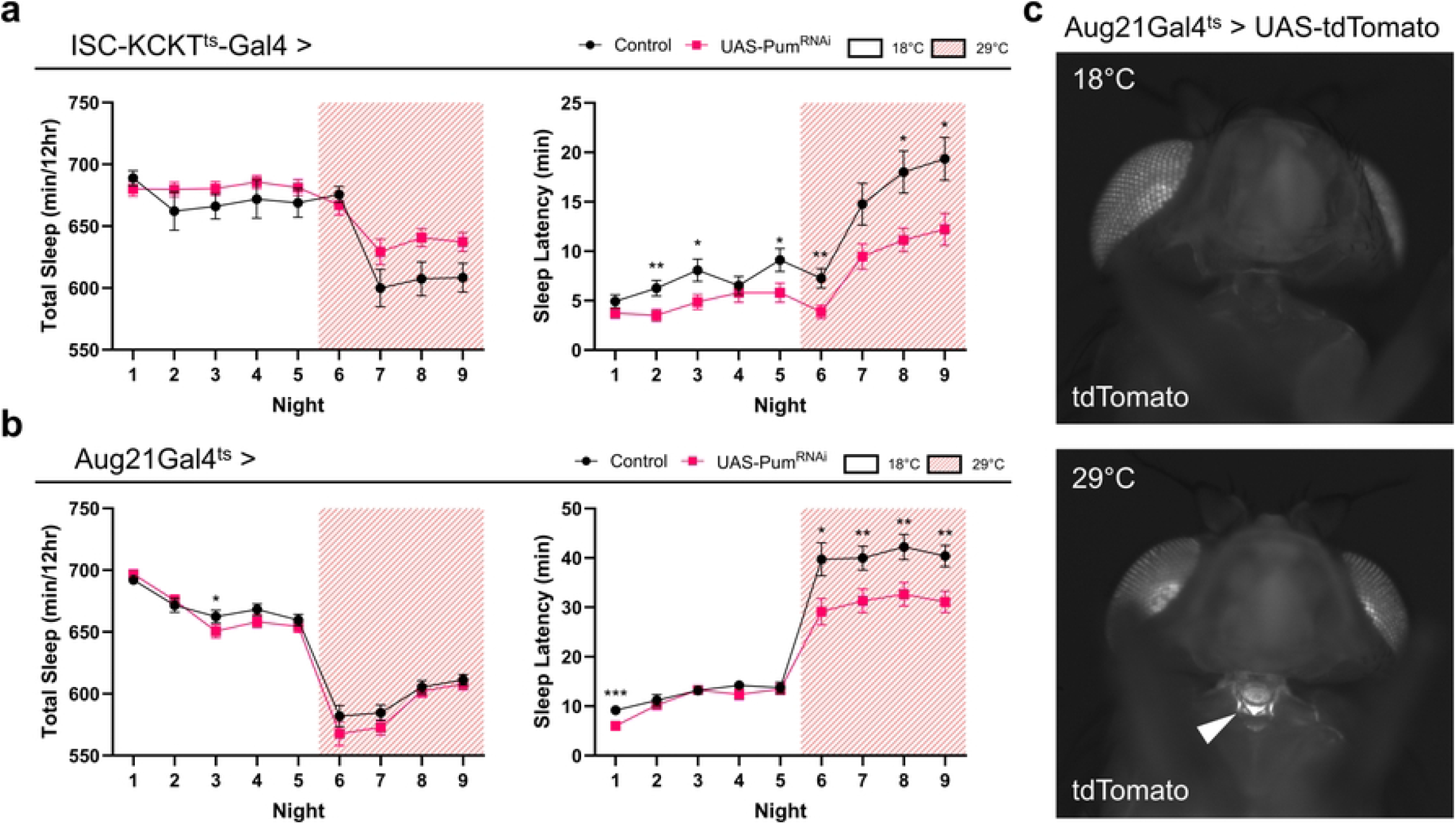
Tissue-specific knockdown of *pumilio* in ISCs or the CA reduces nighttime sleep latency. (a) Nighttime total sleep (left) and sleep latency (right) for male flies with *pum* knockdown specifically in ISCs (*ISC-KCKT*^*ts*^*-Gal4 >UAS-pum*^*RNAi*^ (VALIUM20), n=53) compared to control flies (*ISC-KCKT*^*ts*^*-GAL4 >UAS-mCherry*^*RNAi*^, n=51). While sleep latency is reduced, a slight non-significant difference in total sleep was observed at 29°C. (b) Nighttime total sleep (left) and sleep latency (right) for male flies with *pum* knockdown specifically in the CA (*Aug21Gal4*^*ts*^ *>UAS-pum*^*RNAi*^ (VALIUM20), n=95) compared to control flies (*Aug21Gal4*^*ts*^ *>UAS-mCherry*^*RNAi*^, n=96). Sleep latency is reduced with no overall difference in total sleep. For (a) and (b) values and error bars represent mean ± s.e.m. Asterisks indicate statistically significant differences between genotypes (* *P <*0.05, ** *P <*0.01, *** *P <*0.001, Mann-Whitney test). (c) Representative fluorescence images demonstrating temperature-dependent expression of a reporter (*Aug21Gal4*^*ts*^ *>UAS-tdTomato*) in the *corpus allatum* at 29°C but not at 18°C. Eye pigment autofluorescence is detected in both images. Arrowhead points at CA.

Temperature-dependent expression in the CA only at 29°C was confirmed by generating a fly with the *Aug21-Gal4*^*ts*^ driver and UAS-tdTomato; fluorescence was observed in the CA only at 29°C (Fig 5c).

## Discussion

In this study, we investigated the role of the RNA-binding protein Pumilio in mediating communication between specific peripheral cell populations and the central circuits that control sleep. Pumilio is a highly conserved post-transcriptional regulator known for its critical functions in maintaining various stem cell populations [19] and, within the brain, for modulating sleep homeostasis in response to sleep loss [45]. However, its function in peripheral tissues as a regulator of basal sleep has remained unexplored. We began by examining the expression of *pum* within the adult *Drosophila* intestine, where analysis of snRNA-seq data revealed that *pum* is broadly expressed across various intestinal cell types, with notable enrichment in ISCs, EBs, a subset of ECs, and the cardia. This finding positioned *pum* as a potential candidate influencing communication from the gut to the brain.

Although we initially focused on manipulating *pum* in ISCs using the commonly employed *esg-Gal4; Su(H)Gal80* driver, recent studies have shown that this driver can also express UAS-transgenes in additional tissues, including the CA [41]. We therefore tested the contribution of both the ISCs and CA. We found that conditional knockdown of *pum* simultaneously in ISCs and CA significantly accelerated the onset of nighttime sleep in male flies. This effect appears to be specific, bidirectional, and dependent on *pum* expression level, as conditional overexpression of *pumilio* in the same cell populations produced the opposite phenotype, an increase in sleep latency. In females, this effect was present but weaker and less consistent among experimental biological replicates.

Pumilio proteins are well-characterized post-transcriptional regulators that typically bind to the 3^*′*^ untranslated regions of target mRNAs to repress their translation or promote their degradation [19, 46]. While neuronally expressed *pum* has been implicated in sleep homeostasis following chronic sleep deprivation [45], our results establish a novel role for Pum in ISCs and the CA influencing basal sleep latency.

To explore potential downstream molecular events in the brain that correlate with the altered sleep phenotype, we performed RNA sequencing on heads from flies with ISC and CA *pum* knockdown. This analysis of the head transcriptome revealed 55 DEGs, indicating that peripheral *pum* manipulation leads to molecular changes in the central nervous system. Gene Ontology analysis of these DEGs showed a significant enrichment for terms related to cellular stress, such as “response to heat” and “protein folding.” This is consistent with our observation of a strong upregulation of multiple heat shock protein (Hsp) genes. Hsps are recognized for their neuroprotective functions and can be induced by various stressors, including conditions like sleep deprivation [47, 48]. The heat-shock pathway has also been linked to the distribution of sleep across the day, with differences seen at 18°C vs 29°C [49].

Our head transcriptome analysis also highlighted other widespread physiological adjustments. For instance, several odorant-binding proteins, genes involved in lipid metabolism (e.g., *Lsd-1, Acsx2*), and cytochrome P450s were differentially expressed.

The alteration in *NT5E-2*, an ecto-5^*′*^-nucleotidase involved in purine metabolism, is also noteworthy given the well-established role of adenosine as a key sleep homeostatic factor [50–53]. These diverse transcriptional changes underscore the complex systemic impact of manipulating Pum function in peripheral tissues.

### Bridging peripheral Pumilio function and central sleep control – potential mechanisms

Our tissue-specific knockdown experiments pinpoint both the ISCs and the CA as sites where Pum activity influences sleep latency. Pum function in the CA, the primary endocrine gland for Juvenile Hormone (JH) biosynthesis [54], and in ISCs, which are critical for gut homeostasis and inter-organ communication [22, 55, 56], suggests that hormonal signaling is a likely mediator. Given the roles of these tissues, the JH signaling pathway, which is known to influence sleep and locomotor activity in *Drosophila* [57, 58], emerges as a plausible candidate for mediating Pum’s effects on sleep. Notably, we did not observe a significant change in the expression of the direct JH-response gene Kr-h1 in the head, suggesting that peripheral *pumilio* knockdown does not simply elevate JH levels in the head. Instead, our data may point to a more nuanced mechanism, such as an alteration in the sensitivity of central circuits to existing JH signals.

Among the upregulated Hsps, *Hsp83* is a particularly compelling candidate for linking peripheral Pum function to central sleep regulation. Hsp83 was one of the most significantly upregulated DEGs in the heads of flies with peripheral *pum* knockdown. Beyond its role as a molecular chaperone, Hsp83 is directly implicated in sleep; it is associated with the Gene Ontology category “regulation of circadian sleep/wake cycle, sleep” (GO:0045187) [59–61]. Importantly, Hsp83 mutant flies exhibit dysregulated sleep homeostasis and increased mortality following sleep deprivation [48]. Furthermore, Hsp83 is an established interactor with the JH signaling pathway. It facilitates the nuclear import of the JH receptor Methoprene-tolerant (Met) and is a target of JH-induced phosphorylation, thereby influencing JH intracellular signaling [62, 63]. Wu et al. [64] have shown that adult-specific knockdown of *Met* within the *α*/*β* lobes of the mushroom body, a key sleep regulatory center [65, 66], reduces nighttime sleep in male flies. Thus, the pronounced upregulation of *Hsp83* in the head following peripheral *pum* knockdown could reflect a systemic stress response, or, more directly, it might modulate the brain’s JH signaling by influencing Met activity, perhaps within the mushroom body, thereby impacting sleep circuits.

Pumilio-binding motifs (5^*′*^-UGUANAUA-3^*′*^; where N = A/C/U/G) have been identified in the 3^*′*^ UTR of numerous human [67] and *Drosophila* transcripts [46, 68]. Whether Pumilio directly regulates heat shock–related transcripts including *Hsp83* transcripts remains to be determined.

In summary, our study identifies a novel role for the RNA-binding protein Pumilio in peripheral tissues regulating sleep latency. We propose a model where knockdown of *pumilio* in the ISCs and CA induces a systemic signal that upregulates *Hsp83* in the brain. This, in turn, may modulate JH signaling sensitivity within central sleep circuits, thereby accelerating sleep onset.

### Limitations and future experiments

In this study, we focused on the role of Pum in ISCs and the CA. However, *pum* is expressed in other intestinal and peripheral tissues, and recent work by Weaver et al. has shown that *esg-Gal4* can drive expression in a small subset of neurons in the adult brain, whereas *Su(H)-Gal4* exhibits broader neuronal expression [69]. Because our experiments combined *esg-Gal4* with *Su(H)GBE-Gal80*, we anticipate that if any neurons were affected, this would be limited to a small subset. Nevertheless, confirming the extent of neuronal expression under these conditions will require future anatomical and functional studies.

Because our experiments relied on temperature shifts to induce transgene expression, we cannot fully exclude subtle effects of elevated temperature on sleep architecture or gene expression, despite identical temperature treatments across genotypes. Future experiments using inducible systems independent of temperature (e.g., RU486/GeneSwitch) could help rule out these confounds.

Our transcriptomic analysis was performed on whole heads rather than dissected brains; thus, some transcriptional changes could arise from non-neuronal head tissues such as fat body or sensory organs. Future work isolating brain tissue specifically will refine the molecular targets of peripheral *pum* manipulation.

In addition, the *esg-Gal4; Su(H)Gal80* driver used is active in both ISCs and the CA. Consequently, the observed transcriptional changes in the head likely represent the combined effects of *pum* knockdown in both tissues. Future work employing tissue-specific drivers for RNA sequencing will be essential to disentangle the distinct molecular signals originating from the gut versus the CA. These studies, in addition to direct measurement of JH titers and functional analysis of Hsp83’s role, will be crucial to fully dissect this regulatory network linking peripheral Pumilio function to the central control of sleep.

We also observed clear sex-specific differences in the sleep phenotypes resulting from *pum* manipulation in ISCs and the CA, with males displaying a stronger and more consistent reduction in sleep latency. Sex-specific differences in sleep regulation have been reported in *Drosophila* (reviewed by Asahina et al. [70]), potentially arising from hormonal modulation or dimorphic physiology of peripheral tissues. The work of Miguel-Aliaga and colleagues has shown that ISCs in both sexes respond to JH and express the *Met* receptor, suggesting that sex-dependent hormonal signaling could shape how peripheral Pumilio activity influences sleep circuits [71, 72]. These findings, together with our observation that *pum* knockdown in ISCs and the CA leads to stronger sleep phenotypes in males, suggest that Pumilio might influence or modulate these hormone-sensitive pathways in a sex-specific manner. Because both ISC physiology and CA function can be modulated by JH and other hormones, it is plausible that the weaker phenotype observed in females reflects differences in hormonal sensitivity or endocrine feedback. Future experiments should test this hypothesis by manipulating JH signaling components, such as *Met* or *Kr-h1*, in a sex- and tissue-specific context, and by quantifying JH titers following *pum* perturbation.

Our RNA-seq analysis revealed several differentially expressed odorant-binding and olfactory-related genes in males following peripheral *pum* knockdown. Interestingly, olfactory circuits in male flies are particularly sensitive to JH regulation, which modulates chemosensory-guided behaviors such as courtship and olfactory preference in a sex-dependent manner [70, 73–75]. This raises the possibility that the upregulation of olfactory genes observed in males reflects enhanced JH signaling or altered neuronal sensitivity downstream of peripheral *pum* manipulation. Future studies should test this hypothesis by examining whether JH signaling components within olfactory neurons respond to intestinal or CA-specific *pum* perturbation and whether similar transcriptional responses occur in females.

While our data supports a peripheral-to-central influence of Pumilio, the nature of the signaling route remains to be determined. Possible mechanisms include humoral endocrine factors such as JH or metabolites released from the gut, immune or cytokine-like signals, or visceral neuronal projections connecting the gut and brain. Future studies combining tissue-specific manipulations with circulating metabolite profiling or neuronal tracing could help delineate these pathways.

Altogether, our findings provide a foundation for dissecting how gut–endocrine–brain axis signaling is influenced by a post-transcriptional regulator such as Pumilio. Given the high conservation of Pumilio and its human homologs (PUM1 and PUM2), it will be interesting to explore whether similar peripheral regulatory mechanisms contribute to sleep or circadian modulation in other organisms.

## Acknowledgments

Stocks obtained from the Bloomington Drosophila Stock Center (NIH P40OD018537) were used in this study. We thank Dr. Heinrich Jasper (Buck Institute for Research on Aging, now at Genentech Inc.) for the *esgGal4*^*ts*^ fly line and Dr. Michael Stern (Rice University) for the UAS-Pum fly line. We thank the members of the Agosto laboratory and the Ghezzi laboratory, especially Yilmaz Berk Koru, Aylin Kaya Koru, and Carolina I. Maldonado-Valedón for helping with sleep assay setup.

